# “Increased risk of thrombocytopenia and death in patients with bacteremia caused by high alpha toxin-producing methicillin-resistant *Staphylococcus aureus*”

**DOI:** 10.1101/2021.07.27.454087

**Authors:** Fatimah Alhurayri, Edith Porter, Rachid Douglas-Louis, Emi Minejima, Julianne Bubeck Wardenburg, Annie Wong-Beringer

**Affiliations:** University of Southern California, Los Angeles, CA, USA; California State University, Los Angeles, CA, USA; Washington University, St. Louis, MO; Huntington Memorial Hospital, Pasadena, CA, USA

**Keywords:** Alpha toxin, virulence factors, *Staphylococcus aureus* bacteremia, mortality, platelets, thrombocytopenia

## Abstract

**Background:** Alpha toxin (Hla) is a major virulence factor of *Staphylococcus aureus* that targets platelets but clinical data on Hla pathogenesis in bacteremia (SAB) is limited.

**Objective:** We examined the link between *in vitro* Hla activity and outcome.

**Methods:** Study isolates obtained from 100 patients with SAB (50 survivors; 50 non-survivors) were assessed for *in vitro* Hla production and activity by Western immunoblotting and hemolysis assay, respectively. Relevant demographics, laboratory and clinical data were extracted from patients’ medical records to correlate Hla activity of the infecting isolates with outcome.

**Results:** Hla production strongly correlated with hemolytic activity (*r*_*s*_=0.93) *in vitro*. A trend towards higher hemolytic activity was observed for MRSA compared to MSSA and with high-risk source infection. Significantly higher hemolytic activity was noted for MRSA strains isolated from patients who developed thrombocytopenia (median 52.48 vs 16.55 HU/ml in normal platelet count, *p*=0.012) and from non survivors (median 30.96 vs 14.87 HU/ml in survivors, *p=* 0.014) but hemolytic activity of MSSA strains did not differ between patient groups.

**Conclusions:** *In vitro* Hla activity of *S. aureus* strains obtained from patients with bacteremia may be used to predict risk for thrombocytopenia and death which supports bedside phenotyping and therapeutic targeting in the future.

## Introduction

*Staphylococcus aureus* is a leading cause of bloodstream infection, affecting an estimated 50 in 100,000 people annually with an overall mortality rate of up to 57 % in adults (1). Despite receipt of antibiotic therapy, one in three patients develop persistent bacteremia, which is associated with complications, prolonged hospitalization, and increased risk of death (2). Numerous factors may contribute to the varied outcomes observed in *S. aureus* bacteremia (SAB) including heterogeneity in host immunity and variable expression of virulence factors across clinical *S. aureus* strains (1).

Alpha toxin (Hla), a water-soluble, 34 kDa monomer, is a well-characterized cytolysin that is known to play a key role in the pathogenesis of *S. aureus* infections. Upon binding of Hla to its host cell receptor, ADAM10 (A Disintegrin And Metalloproteinase domain-containing protein-10) which is widely expressed on endothelial cells, epithelial cells, and immune cells, oligomerization of the toxin occurs leading to heptameric pores in the membranes and subsequent alterations in cellular signaling and cellular lysis (3). Platelets are the most abundant, non-nucleated immune cells in circulation and have been shown to kill *S. aureus* directly and indirectly through functional enhancement of other immune cells such as macrophages (4). Wuescher et al found reduced survival rate with increased cytokine storm and higher bacterial load in kidneys of platelet-depleted mice infected with USA300 causing bacteremia compared to wild-type (WT) mice (5). Importantly, recent studies have demonstrated that Hla targets platelets and causes platelet activation, aberrant aggregation, and injury (6, 7). Surewaard et al found more thrombocytopenia and significantly higher platelet aggregation in the livers of mice that were infected with wild-type *S. aureus* compared to an isogenic *hla*-deletion mutant (8).

Additionally, Hla was shown to induce platelet desialylation resulting in enhanced and premature platelet clearance through the hepatic Ashwell-Morell receptor (AMR) (7). High Hla production was shown to increase severity and reduce survival of infection in numerous experimental models of pneumonia (9, 10), sepsis (6), peritonitis (11), and brain abscesses (12). Administration of a novel anti-alpha toxin monoclonal antibody developed to specifically target and neutralize Hla was shown to provide survival benefit in multiple animal models of infection, including pneumonia (13, 14), skin and soft-tissue infections (15, 16), and bacteremia (17). Despite the extensive research that highlights Hla as a major toxin in animal models and the considerable efforts towards developing antibodies that target and neutralize Hla in *S. aureus* infections, few studies assessed the relationship between *in vitro* Hla production by clinical isolates as a possible predictive marker for the development of thrombocytopenia and patient outcomes in SAB. Given the previously reported variation in Hla expression across *S. aureus* strains (18, 19) and its involvement in the pathogenesis of *S. aureus* infections, we hypothesized that *S. aureus* bloodstream isolates produce varying Hla levels and that high Hla-producing strains are associated with increased risk of thrombocytopenia and mortality in patients with bacteremia. Our study objectives were to: 1) measure the *in vitro* level of Hla production and hemolytic activity of *S. aureus* bloodstream isolates and 2) correlate Hla activity with platelet count and outcome of SAB.

## Methods

### Patient and bacterial isolate selection

Study isolates and clinical data had been previously collected as part of a large multicenter prospective observational study of adult patients hospitalized for SAB from two affiliated medical centers in Los Angeles, USA. The study was approved by the respective IRB at each site; informed consent was waived as the study was observational in design. A total of 100 study isolates were selected to represent equal numbers of patients who died or survived within 30 days of bacteremia onset. Survivors were those who had favorable outcomes: bacterial clearance and clinical improvement by day 4 following onset of SAB and end of therapy success while non-survivors were those whose death was deemed SAB-related in the patient’s medical record and had persistent bacteremia or lack of clinical improvement or worsening on day 4 following onset of SAB. Thrombocytopenia was defined as platelet count < 150 × 10^9^/L at time of SAB onset. Source of infection was grouped based on risk of mortality as previously defined: high (> 20%; endovascular, lower respiratory tract, intrabdominal, and central nervous system foci), intermediate (10–20%; osteoarticular, soft tissue, and unknown sources), and low (< 10%; IV catheter, urinary tract infection, ear-nose-larynx, gynecologic sources, and several manipulation-related sources including digestive endoscopy, arterial catheterization, and sclerosis of esophageal varices) (20).

Patients’ medical records were retrospectively reviewed to extract relevant demographics, laboratory and clinical data (see **Table 1**), and managed using REDCap electronic data capture tools hosted at University of Southern California (21). Our *in vitro* analysis included 100 clinical isolates from those patients plus three *S. aureus* control strains: LAC (USA300) *hla* wild type, isogenic *Δhla* mutant lacking Hla production, and *Δhla*-complemented mutant strains with restored Hla production (22).

**Table 1.**
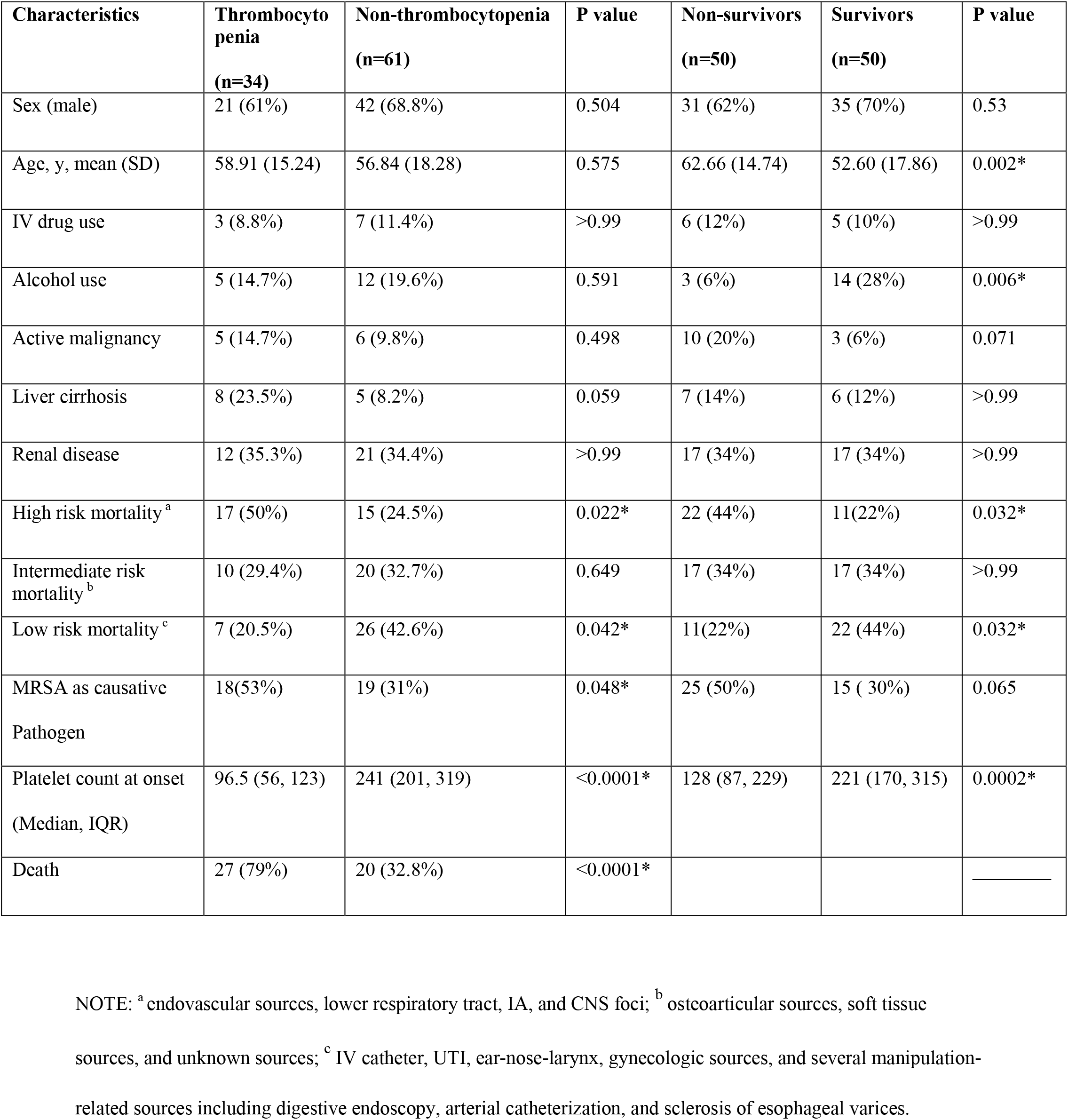
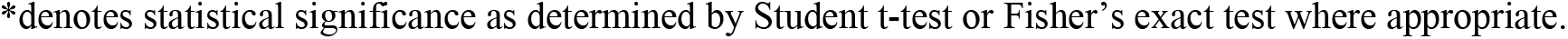
Comparison of patient characteristics grouped by development of thrombocytopenia and 30-day survival.

| Characteristics | Thrombocyto<br>penia<br>(n=34) | Non-thrombocytopenia<br>(n=61) | P value | Non-survivors<br>(n=50) | Survivors<br>(n=50) | P value |
| --- | --- | --- | --- | --- | --- | --- |
| Sex (male) | 21 (61%) | 42 (68.8%) | 0.504 | 31 (62%) | 35 (70%) | 0.53 |
| Age, y, mean (SD) | 58.91 (15.24) | 56.84 (18.28) | 0.575 | 62.66 (14.74) | 52.60 (17.86) | 0.002* |
| IV drug use | 3 (8.8%) | 7 (11.4%) | >0.99 | 6 (12%) | 5 (10%) | >0.99 |
| Alcohol use | 5 (14.7%) | 12 (19.6%) | 0.591 | 3 (6%) | 14 (28%) | 0.006* |
| Active malignancy | 5 (14.7%) | 6 (9.8%) | 0.498 | 10 (20%) | 3 (6%) | 0.071 |
| Liver cirrhosis | 8 (23.5%) | 5 (8.2%) | 0.059 | 7 (14%) | 6 (12%) | >0.99 |
| Renal disease | 12 (35.3%) | 21 (34.4%) | >0.99 | 17 (34%) | 17 (34%) | >0.99 |
| High risk mortality<sup>a</sup> | 17 (50%) | 15 (24.5%) | 0.022* | 22 (44%) | 11(22%) | 0.032* |
| Intermediate risk<br>mortality<sup>b</sup> | 10 (29.4%) | 20 (32.7%) | 0.649 | 17 (34%) | 17 (34%) | >0.99 |
| Low risk mortality<sup>c</sup> | 7 (20.5%) | 26 (42.6%) | 0.042* | 11(22%) | 22 (44%) | 0.032* |
| MRSA as causative<br>Pathogen | 18(53%) | 19 (31%) | 0.048* | 25 (50%) | 15 ( 30%) | 0.065 |
| Platelet count at onset<br>(Median, IQR) | 96.5 (56, 123) | 241 (201, 319) | <0.0001* | 128 (87, 229) | 221 (170, 315) | 0.0002* |
| Death | 27 (79%) | 20 (32.8%) | <0.0001* |  |  | _____ |

NOTE: <sup>a</sup> endovascular sources, lower respiratory tract, IA, and CNS foci; <sup>b</sup> osteoarticular sources, soft tissue sources, and unknown sources; <sup>c</sup> IV catheter, UTI, ear-nose-larynx, gynecologic sources, and several manipulation- related sources including digestive endoscopy, arterial catheterization, and sclerosis of esophageal varices.
\*denotes statistical significance as determined by Student t-test or Fisher's exact test where appropriate.

### *In vitro* measurement of Hla expression

A single colony of each study isolate freshly grown on a TSA plate was inoculated in 5 ml Tryptic Soy Broth and incubated overnight at 37°C with shaking at 200 rpm. The OD_600nm_ was adjusted to 0.1 for all isolates and the bacterial suspension was incubated at 37°C for another 20 h to reach stationary growth phase (23, 24). Then, an aliquot of the culture was removed for CFU determination by plate counting and the remainder was centrifuged at 4°C and 3,100 rpm (Eppendorf Centrifuge 5415R) for 10 min to generate cell-free supernatants, which were stored at −80°C until later analysis. All cell-free supernatants were normalized to 1×10^9^ CFU and used in both Western immunoblot and hemolysis assays.

#### Western immunoblot assay

Ten μL of cell-free culture supernatants were subjected to SDS-PAGE using 4-20% Tris-Glycine extended gels. Purified Hla (Sigma-Aldrich, St. Louis, US) was used as controls diluted to in 250 ng, 125 ng, and 25ng/10 μl samples. Proteins were then transferred to a PVDF membrane and blocked with 5% BSA in TBST (Tris-buffered saline, 0.1% Tween 20, TBST) for 2 h, then incubated overnight at 4**°**C with the primary antibody mouse anti-SA Hla [8B7] -N-terminal (Abcam, Cambridge, United Kingdom) diluted 1:2500 in the blocking buffer. After three five-min TBST washes, the blot was incubated with goat-anti Mouse-HRP (Abcam, US) as secondary antibody diluted 1:2000 in the blocking buffer for 1 h followed by a 5 min wash in TBST. The signal was developed using TMB (3,3’,5,5’-tetramethylbenzidine) Peroxidase (HRP) Substrate Kit according to the manufacturer’s instructions (Vector kit SK4400, Vector Laboratories, Burlingame, US). Developed membranes were dried and images were acquired with a Bio-Rad ChemiDoc Touch Gel Imager (Bio-Rad, Hercules, US) and the quantitative analysis was performed using Image Lab software (6.0.1 version).

#### Hemolysis assay

The hemolytic activity of Hla was evaluated by measuring the hemolysis of rabbit erythrocytes (Innovative research, US). As described previously (25), 100 µl of serially diluted (1:5 up to 1:640 in PBS) culture supernatant was added to a round-bottom polysterene 96-well plate followed by the addition of 100 µl of washed rabbit erythrocytes (1% in PBS). The plate was incubated for 1 h at 37**°**C and then centrifuged at room temperature, 220 rpm for 10 minutes. One hundred µl from each well was transferred to a new flat-bottom polysterene 96-well plate and absorbance was measured at 570 nm. PBS was used as a negative control and 5000 ng purified Hla resuspended in 100 μl PBS was used as a positive control. Hla level (in hemolytic units per ml, HU/ml) was defined as the inverse of the dilution causing 50% hemolysis. All supernatants were tested in duplicates and the results were averaged.

### Data Analysis

The correlation between Hla protein level measured by Western immunoblotting and the hemolytic activity was examined for 61 clinical isolates representing 40 MRSA; 21 MSSA; survivors and non-survivors. The relationship between the hemolytic activity of all 100 SAB isolates relative to methicillin resistance and platelet count at onset of bacteremia, source risk of infection, and 30-day mortality was analyzed.

Statistical analysis was performed using GraphPad Prism (version 9.0). Unpaired t-test, Mann-Whitney tests and Fisher’s Exact tests were used to test continuous and categorical variables where appropriate. Spearman correlation test was performed for correlation analysis. A two-tailed p-value of < 0.05 denotes statistical significance.

## Results

### Patient characteristics and platelet trends

**Table 1** compares the characteristics between those with and without thrombocytopenia and between survivors and non-survivors in terms of sex, age, IV drug use, and comorbid conditions that may predispose to the development of thrombocytopenia, cause and source risk of infection, and platelet count at onset. Thrombocytopenia occurred in 36 % (34/95) of our study cohort overall, with 8-times greater proportion among non-survivors than survivors (57 %, 27/47 vs 14%, 7/48; p<0.0001, OR 7.91, CI: 2.89 to 19.44). Among those who developed thrombocytopenia at onset of SAB, a greater proportion had sources of infection associated with high risk for death such as endovascular and lower respiratory tract infections (50 % vs 24 %, p=0.022) and had MRSA as a causative pathogen (53% vs 31%, p=0.048) when compared to those who did not, though the groups did not differ in age or comorbid conditions except for liver cirrhosis (24% vs 8%, p=0.059). On the other hand, non-survivors were older (median 62 vs. 52 years, p=0.002) with a trend towards greater proportion with active malignancy (20 % vs 6 %, p=0.071). Similarly, non-survivors were also more likely to have high risk source infections (44 % vs 22 %, p=0.032).

Of the 95 patients with available platelet counts at onset of SAB, the median platelet counts were 96.5, 241, 221, 128 × 10^9^/L in the following groups respectively: thrombocytopenic, non-thrombocytopenic, survivors, and non-survivors. An analysis of platelet dynamics during the first 7 days of bacteremia showed a decline of platelet counts through day 4 with negligible recovery among non-survivors relative to survivors (median platelet count, 1 × 10^9^/L): on day 1 [128 (IQR 87, 229) vs 221 (IQR 170, 315); p=0.0002], on day 4 [83 (IQR 51,159 vs 200 (IQR 161, 292); p<0.0001), and on day 7 [91 (IQR 44.7, 163) vs 237 (IQR 189, 369); p <0.0001) (**Figure 1**).

**Figure 1.**
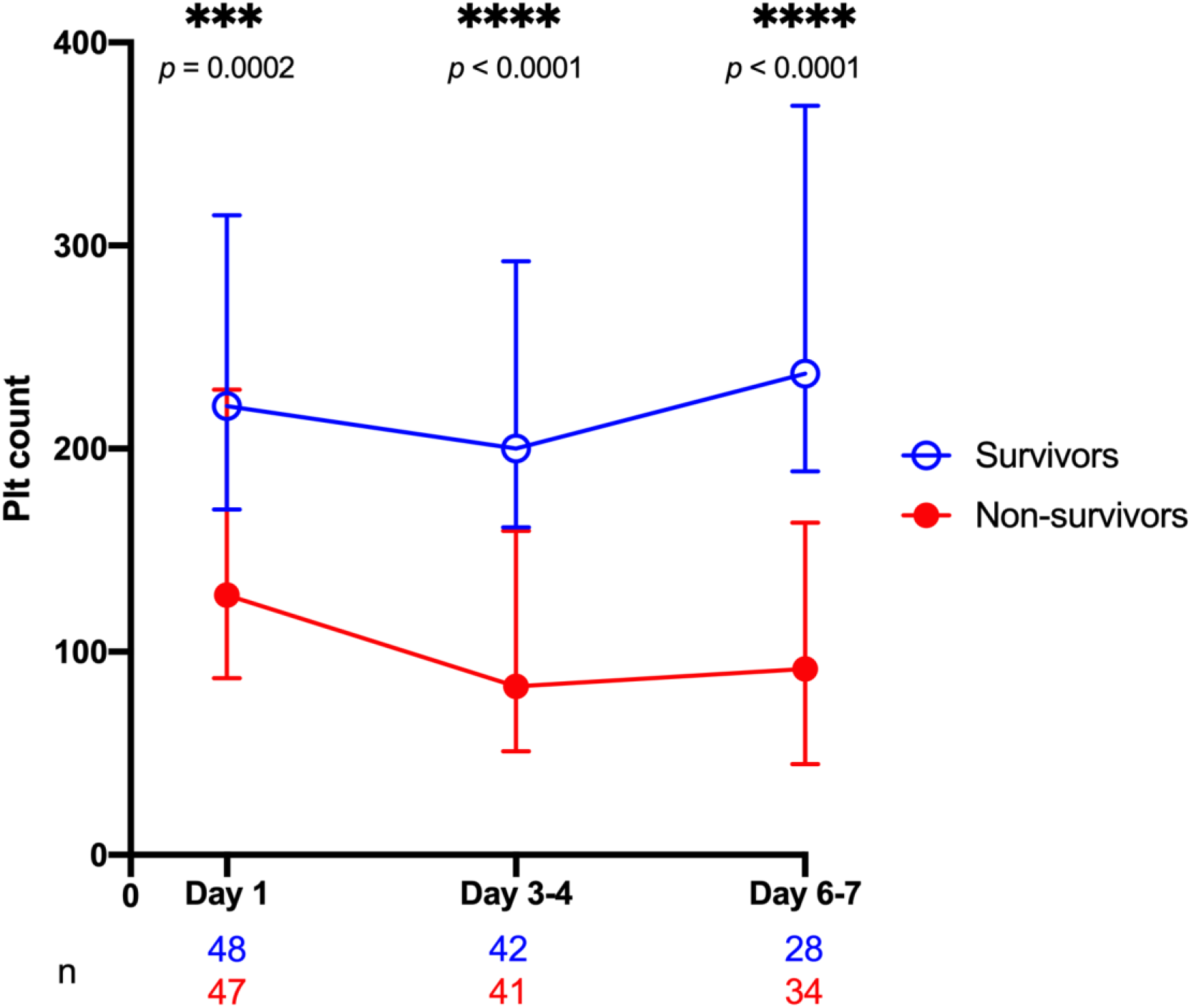
Association between platelets dynamics during SAB. Strong positive association between serial measurements of platelets (plt) counts during first 7 days of bacteremia and survival. *p* values as determined by Mann-Whitney tests.

### High correlation of Hla protein level and hemolytic activity

We evaluated Hla expression by first determining the correlation between protein levels and hemolytic activity measured by Western immunoblot and hemolysis assays, respectively, in a subset of 61 clinical isolates, representing survivors and non-survivors, MRSA and MSSA isolates, plus the three control strains (**Figure 2**). A very strong correlation was found between Hla protein level and hemolytic activity (*r*_*s*_= 0.93, *p* <0.0001). Based on these findings and considering the labor intensity of the Western immunoblot, we performed hemolysis assays only on the remaining clinical isolates and all subsequent analysis was based on results obtained from the hemolysis assay.

**Figure 2.**
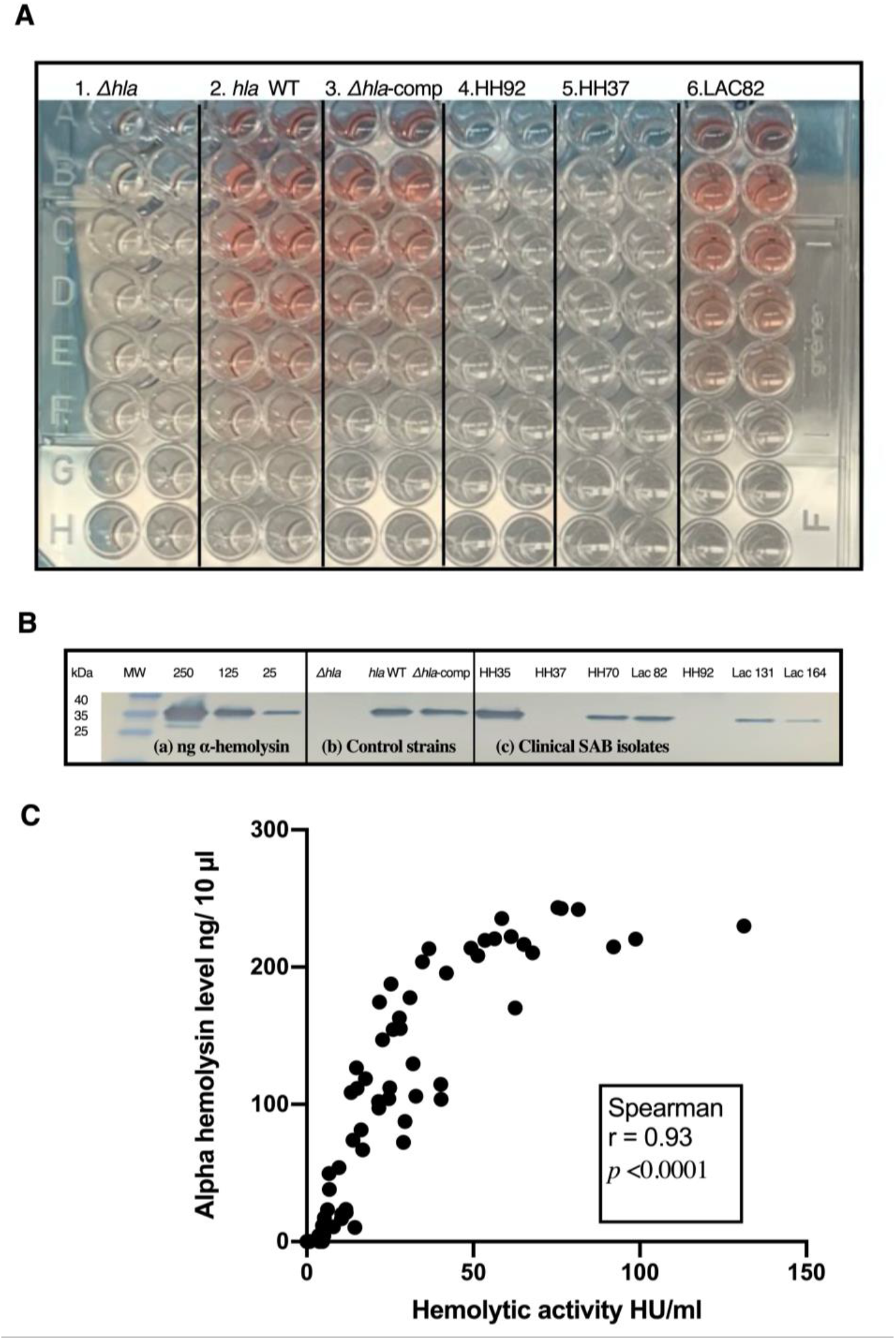
Correlation of Western immunoblot and hemolysis assays. **(A)** Representative image of hemolysis assay with supernatants that were tested in duplicates, collected from control strains (1. isogenic *Δhla* mutant lacking Hla production, 2. LAC (USA300) *hla* wild type and 3. *Δhla*-complemented mutant strains with restored Hla production) and clinical isolates (4. HH92, 5. HH37, and 6. LAC82) in a 96 well plate. **(B)** Western immunoblot for (a) known amounts of purified Hla, (b) control strains and (c) representative SAB clinical isolates. **(C)** Correlation between Hla protein concentration and hemolytic activity, *p* values as determined by Spearman correlation test.

### Association of high hemolytic activity with thrombocytopenia and mortality

Bloodstream *S. aureus* isolates exhibited a wide variation of Hla expression in terms of hemolytic activity (range: 0 to 138.7 HU/ml). Overall, higher hemolytic activity was observed for MRSA vs MSSA isolates (25.12 vs 14.76 HU/ml, *p*= 0.09), isolates from patients with thrombocytopenia vs those from patients with normal platelet count [median 28.1 HU (IQR 7.62, 61.1) vs 16.34 HU (IQR 8.8, 32.3); *p=* 0.08], and from survivors vs non-survivors [median 23.1 HU (IQR 7.62, 59.2) vs 17.97 HU (IQR 5.42, 33.73); *p=* 0.252], respectively (**Figures 3A-B, 4A-B**). When isolates were grouped based on methicillin resistance, a striking difference in hemolytic activity was observed for MRSA isolates from patients with thrombocytopenia vs normal platelet count [median 52.48 HU (IQR 22.19, 69.71) vs 16.55 HU (IQR 9.8, 26.82), *p* = 0.011] and from patients who died vs those who survived [median 30.96 HU/ml (IQR 17, 66.50) vs 14.87 HU/ml (IQR 3.76, 29.53), *p=* 0.014) (**Figures 3C and 4C**).

**Figure 3.**
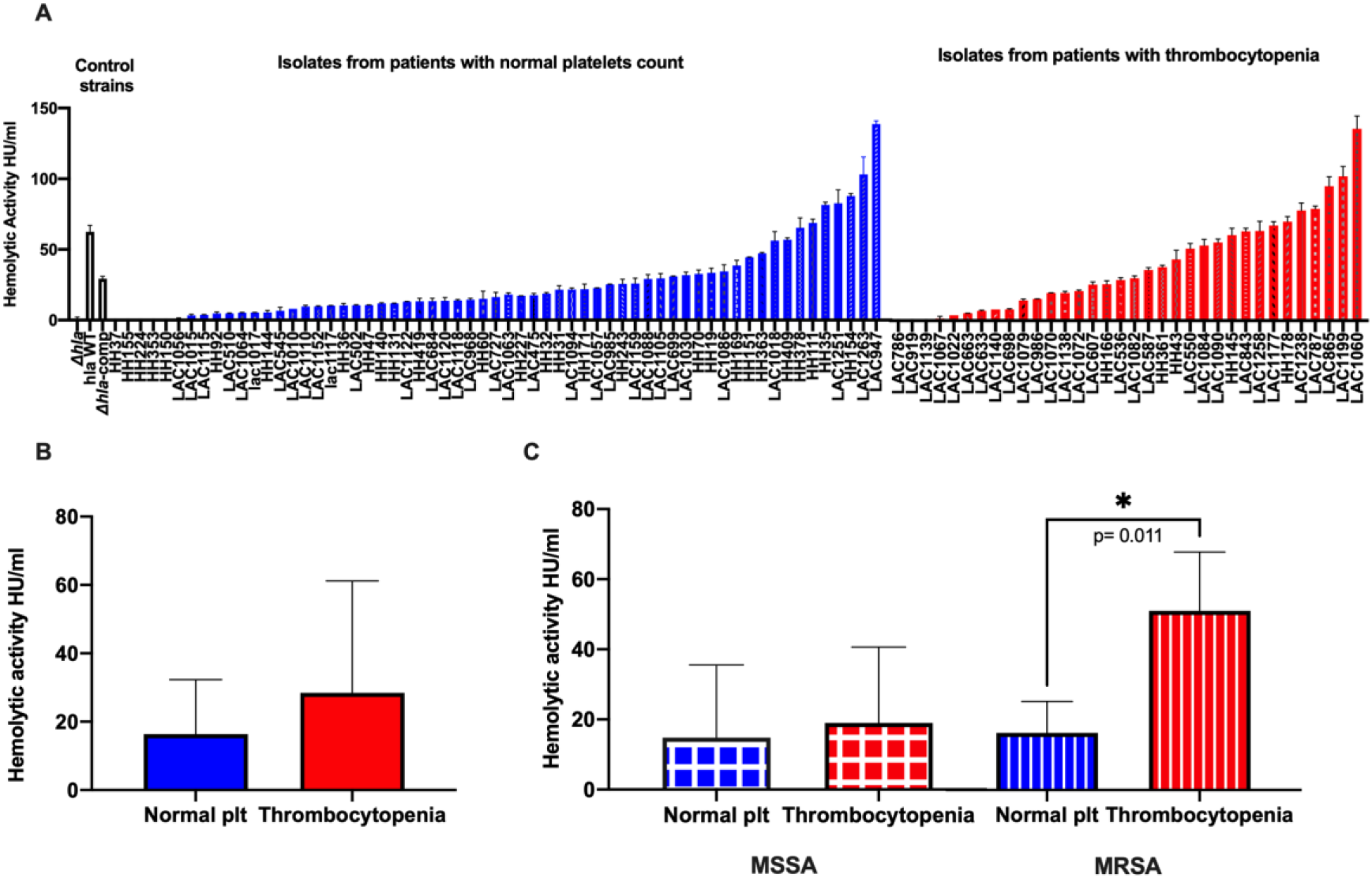
**(A)** Distribution of hemolytic activity across SAB isolates grouped by platelet status. Control strains: LAC (USA300) *hla* wild type, isogenic *Δhla* mutant lacking Hla production, and *Δhla*-complemented mutant strains with restored Hla production. **(B)** Comparison of *S. aureus* hemolytic activity from patients with normal platelet count and thrombocytopenia (including both MSSA and MRSA). **(C)** Comparison of *S. aureus* hemolytic activity from patients with normal platelet count and thrombocytopenia grouped by MSSA or MRSA. plt: platelet count. p values as determined by Mann-Whitney test.

On the contrary, MSSA isolates exhibited similar Hla activity regardless of platelet count status of the infected patients [median 14.76 HU (IQR 6.4, 35.5) vs 18.9 HU (IQR 5.7, 40.6); *p* = 0.91] (**Figure 3C**) or survival status [median 18.99 HU (IQR 5.45, 34.50) non-survivors vs 13 HU (IQR 4.91, 39.32) survivors, *p*= 0.353] (**Figure 4C**). Notably, the mean platelet count among MRSA-infected group was significantly lower compared to MSSA group [155.8 +/- 87.98 SD vs 237 +/- 125.6 SD, *p* = 0.0008]. These findings support the association between Hla activity, thrombocytopenia, and death among MRSA but not MSSA bloodstream isolates.

**Figure 4.**
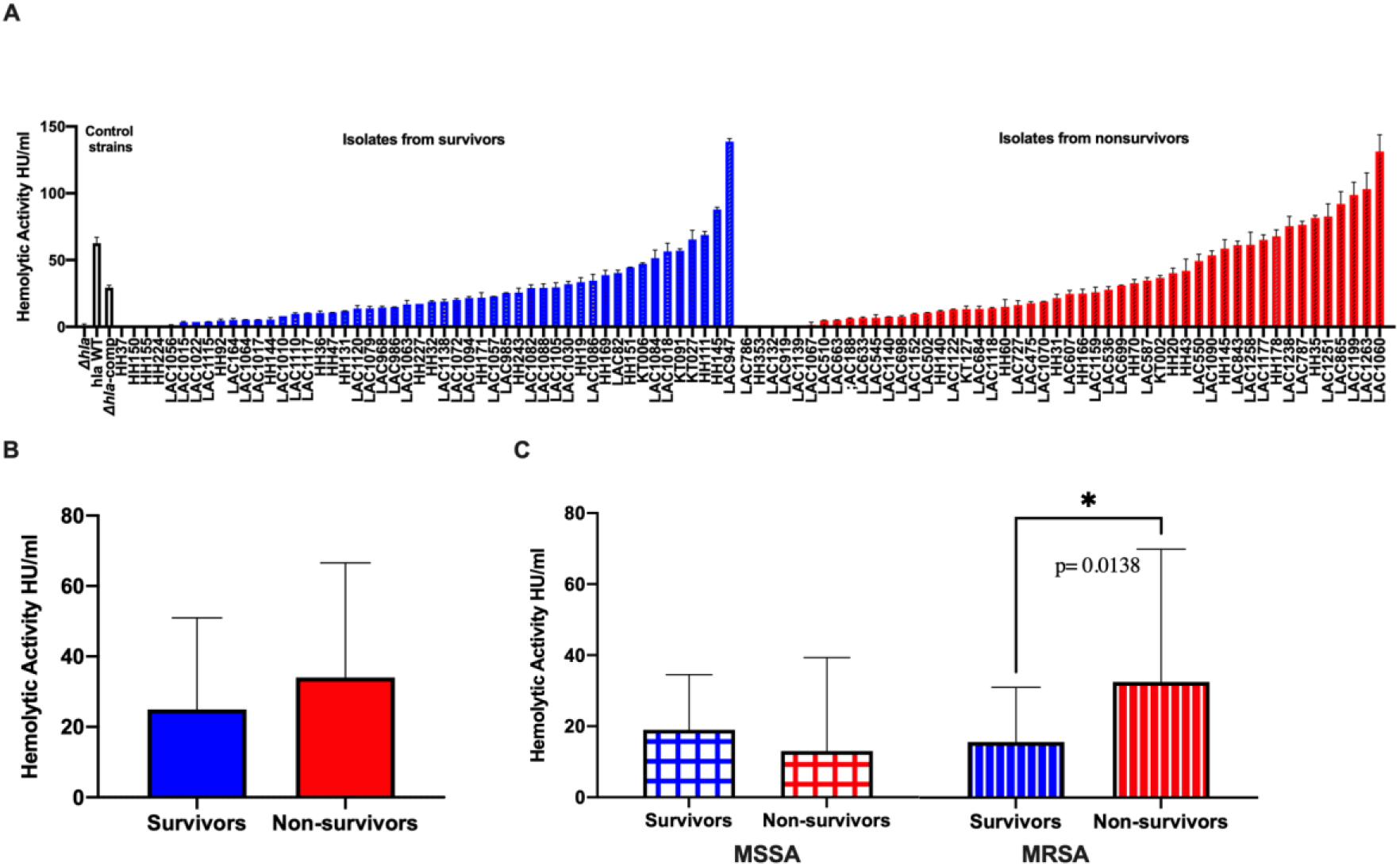
**(A)** Distribution of alpha hemolysin hemolytic activity across SAB isolates grouped by survival status. Control strains: LAC (USA300) *hla* wild type, isogenic *Δhla* mutant lacking Hla production, and *Δhla*-complemented mutant strains with restored Hla production. **(B)** Comparison of *S. aureus* hemolytic activity from survivors and non-survivors (including both MSSA and MRSA). **(C)** Comparison of *S. aureus* bloodstream isolates hemolytic activity from survivors and non-survivors grouped by MSSA or MRSA. p values as determined by Mann-Whitney test.

### Association of high hemolytic activity with high-risk source of infection

We grouped the study isolates according to the source of infection relative to the associated risk of death as reported by Soriano et al (20). We found a positive trend when correlating hemolytic activity with risk source of infection [median 29.04 HU (IQR 12.08, 59.84) vs 15.06 HU (IQR 6.25, 34.74), p=0.066] comparing between isolates from high-risk source (n=33) *versus* intermediate-risk and low-risk sources of bacteremia (n=67), respectively, but this trend did not reach statistical significance (data not shown).

## Discussion

The objective of our study was to measure the *in vitro* level of Hla production and hemolytic activity of *S. aureus* bloodstream isolates and investigate the association of *S. aureus* Hla activity with platelet count and outcome in patients with SAB. We examined a total of 100 *S. aureus* isolates that caused bacteremia chosen to represent equal number of patients who died or survived. From these isolates, *in vitro* Hla production and hemolytic activity were measured by Western immunoblotting and rabbit erythrocyte-based hemolysis assays using cell-free culture supernatants obtained from bacteria in stationary growth phase. We demonstrated an excellent correlation between hemolysin production *in vitro* and hemolytic activity. Consistent with previous reports (26, 19), our findings showed a wide variation in hemolytic activity across bloodstream isolates.

A strong association between thrombocytopenia, defined as a platelet count less than 150 × 10^9^/L, and 30-day mortality has been previously shown by a retrospective study involving 1052 patients with SAB (27). In a separate study that included 49 patients with SAB, strains with high Hla expression were associated with thrombocytopenia (platelet count < 100 × 10^9^/L) from the initial blood sample and death in 4 of 9 patients (7). We extended these findings by analyzing the association of hemolytic activity of the SAB strains and platelet count measured at multiple time points during SAB as well as mortality in a larger patient cohort.

Notably, we showed a strong association between thrombocytopenia and 30-day mortality and found that platelet count nadir occurred around day 4 from onset of bacteremia. These findings provide clear support of the current clinical practice that considers low platelet count as a poor prognostic indicator in SAB, indicative that platelet count should be followed serially particularly during the early course of bacteremia in line with previous studies that analyzed the association between mortality and multiple time points of platelet count in critically ill patients (28, 29).

Previous studies provided mechanistic insights into Hla-mediated platelet dysfunction and depletion in experimental models of *S. aureus* sepsis (6, 8). Therefore, we analyzed the relationship between Hla expression of the infecting *S. aureus* isolates and thrombocytopenia and death in patients with bacteremia. Importantly, our results confirmed the relationship between Hla expression and thrombocytopenia and cast a new light on the contribution of methicillin resistance. We found that MRSA bloodstream isolates had higher overall hemolytic activity and that high Hla-producing MRSA strains are significantly associated with thrombocytopenia and death in *S. aureus* bacteremia but the association between high hemolytic activity and poor outcome was not observed with MSSA strains. In line with these results, Coia et al. measured Hla production in a total of 201 isolates of MRSA and MSSA obtained from various body sites and reported significantly higher Hla production *in vitro* in MRSA relative to MSSA isolates (30). Our results are in partial agreement with Jacobsson et al. as they found a positive association between Hla hemolytic activity (assessed by measuring the zones of hemolysis in nutrient agar plates using rabbit erythrocytes) and complicated bacteremia, but not mortality (31). On the other hand, Sharma-Kuinkel et al reported an inverse association between SAB patients’ outcome and the *in vitro* Hla production (measured by Western blot and ELISA) and hemolytic activity (measured by rabbit erythrocyte lysis assay) (25). One potential explanation is the difference in the patient population studied since their population was limited to postsurgical patients and those on hemodialysis whereby our study included a broader range of patient population. Importantly, Sharma-Kuinkel et al did not analyze their results relative to methicillin resistance which could potentially mask differences between MRSA and MSSA with respect to Hla activity in association with patient outcome as was noted in our study.

Epidemiologic data from a recent retrospective analysis of 92,089 patients indicated that bacteremia caused by MRSA was associated with longer hospitalization, higher rate of readmission with bacteremia recurrence, and increased mortality compared to MSSA (32). Two previous meta-analyses yielded similar findings when comparing mortality rates between patients infected with MRSA and MSSA (33, 34). Our findings suggest differences in virulence of the infecting strains and the Hla-mediated injury to platelets may offer a biologically plausible explanation for the observed difference in mortality between MRSA and MSSA bacteremia. It is possible that MRSA and MSSA bloodstream isolates in our cohort differ in genetic backgrounds and that additional virulence factors may be present in MRSA strains that act synergistically with Hla, thereby contributing to the observed difference in patient outcome. Another potential explanation for the difference in hemolytic activity in our MRSA and MSSA strains is the variation between MRSA vs. MSSA in the regulatory component, *agr* system. Agr (the accessory gene regulator) is known to regulate the expression of many virulence genes including Hla (9) and Cheung et al. reported that *agr* enhanced the regulation of methicillin resistance genes (*mecA, mecR1*) in community-associated MRSA (CA-MRSA) (35). Of interest, Otto et al. have previously linked the possibility of increased virulence of CA-MRSA with the increased expression of *hla*, partially because of the enhanced activity of Agr (36). Moreover, high Hla protein was detected among highly virulent ST93 CA-MRSA strains while *agr*-deficient strains displayed decreased *hla* expression (37). Similarly, others have shown increase in the expression of *hla* and *agr* in ST59 CA-MRSA isolates (38).

Our study on patients with SAB corroborated published literature on the deleterious effect of *S. aureus* Hla on platelets (e.g. platelet aberrant aggregation and desialylation) *in vitro* and in experimental models of sepsis (6, 7) and lend support for future investigations on measuring the virulence phenotype of the infecting strain and therapeutically targeting Hla-mediated effects on platelets to improve patient outcome. Specifically, future studies should examine the feasibility of performing phenotypic assays to measure Hla expression of the infecting *S. aureus* isolates that could be readily adopted into the routine workflow in clinical microbiology laboratory at the time of organism identification. As platelet count measurements are part of routine complete blood count, clinicians should closely monitor platelet count especially during the initial 4 days following onset of *S. aureus* bacteremia relative to Hla expression in the infected strains. Additionally, *in vivo* studies confirming the modulatory potential of existing therapeutics including antibiotics with antivirulence activity (39) on Hla expression and their benefit in mitigating Hla-mediated platelet dysfunction and depletion could accelerate the translation of our findings to practice.

We acknowledge several limitations in our study. First, we measured the *in vitro* Hla production of *S. aureus* bloodstream isolates and correlated it with patient outcome; however, it remains unclear how well *in vitro* Hla production and hemolytic activity correlates with *in vivo* production during bacteremia. In murine models of pneumonia and skin and soft tissue infection, Berube et al showed a direct correlation between *in vivo* Hla production and the degree of tissue injury(40). Nonetheless, the harmful effects of Hla on platelets are supported by a clear relationship observed between high hemolytic activity of the infecting strains and thrombocytopenia during bacteremia in humans. Furthermore, it is possible that other staphylococcal cytotoxins including the beta, delta, and gamma hemolysins and the bi-component leukocidins may have contributed collectively to the observed *in vitro* hemolytic activity, as their role in the pathogenesis of *S. aureus* diseases has not been fully clarified (41, 42). Finally, we observed very strong signals of high Hla proteins in the Western immunoblot for some of the strains which suggest that the measurements may have reached or exceeded the level of saturation. Therefore, we employed the standard rabbit erythrocyte-based hemolysis assay to capture the variable expression of Hla across our bloodstream isolates. The lack of hemolysis exhibited by our *hla*-deletion strain and the recovery of hemolytic activity of the complemented mutant strain support the validity of the hemolysis assay.

In conclusion, our findings show that patients with MRSA bacteremia are more likely to be infected with high Hla-producing strains and are at higher risk for developing thrombocytopenia and death. Importantly, our study provides support for the future *precision* treatment of infections by phenotyping pathogen virulence and host platelet response to stratify patients who may benefit from treatment that target the Hla-platelet interface thereby improving outcome of *S. aureus* bacteremia.

## Acknowledgments

We thank Anne Au, the Clinical Laboratory staff at Huntington Hospital and Los Angeles County-University of Southern California Medical Center for assistance in collection of bacterial specimens.

## Author Contributions

Rachid Douglas-Louis and Emi Minejima contributed by collecting and providing the clinical data.

## References

1. van Hal SJ, Jensen SO, Vaska VL, Espedido BA, Paterson DL, Gosbell IB. 2012. Predictors of mortality in Staphylococcus aureus Bacteremia. Clin Microbiol Rev 25:362–86.

2. Minejima E, Bensman J, She RC, Mack WJ, Tuan Tran M, Ny P, Lou M, Yamaki J, Nieberg P, Ho J, Wong-Beringer A. 2016. A Dysregulated Balance of Proinflammatory and Anti-Inflammatory Host Cytokine Response Early During Therapy Predicts Persistence and Mortality in Staphylococcus aureus Bacteremia. Crit Care Med 44:671–9.

3. Berube BJ, Bubeck Wardenburg J. 2013. Staphylococcus aureus alpha-toxin: nearly a century of intrigue. Toxins (Basel) 5:1140–66.

4. Ali RA, Wuescher LM, Dona KR, Worth RG. 2017. Platelets Mediate Host Defense against Staphylococcus aureus through Direct Bactericidal Activity and by Enhancing Macrophage Activities. J Immunol 198:344–351.

5. Wuescher LM, Takashima A, Worth RG. 2015. A novel conditional platelet depletion mouse model reveals the importance of platelets in protection against Staphylococcus aureus bacteremia. J Thromb Haemost 13:303–13.

6. Powers ME, Becker RE, Sailer A, Turner JR, Bubeck Wardenburg J. 2015. Synergistic Action of Staphylococcus aureus alpha-Toxin on Platelets and Myeloid Lineage Cells Contributes to Lethal Sepsis. Cell Host Microbe 17:775–87.

7. Sun J, Uchiyama S, Olson J, Morodomi Y, Cornax I, Ando N, Kohno Y, Kyaw MMT, Aguilar B, Haste NM, Kanaji S, Kanaji T, Rose WE, Sakoulas G, Marth JD, Nizet V. 2021. Repurposed drugs block toxin-driven platelet clearance by the hepatic Ashwell-Morell receptor to clear Staphylococcus aureus bacteremia. Sci Transl Med 13.

8. Surewaard BGJ, Thanabalasuriar A, Zeng Z, Tkaczyk C, Cohen TS, Bardoel BW, Jorch SK, Deppermann C, Bubeck Wardenburg J, Davis RP, Jenne CN, Stover KC, Sellman BR, Kubes P. 2018. alpha-Toxin Induces Platelet Aggregation and Liver Injury during Staphylococcus aureus Sepsis. Cell Host Microbe 24:271–284 e3.

9. Bubeck Wardenburg J, Patel RJ, Schneewind O. 2007. Surface proteins and exotoxins are required for the pathogenesis of Staphylococcus aureus pneumonia. Infect Immun 75:1040–4.

10. Bartlett AH, Foster TJ, Hayashida A, Park PW. 2008. Alpha-toxin facilitates the generation of CXC chemokine gradients and stimulates neutrophil homing in Staphylococcus aureus pneumonia. J Infect Dis 198:1529–35.

11. Rauch S, DeDent AC, Kim HK, Bubeck Wardenburg J, Missiakas DM, Schneewind O. 2012. Abscess formation and alpha-hemolysin induced toxicity in a mouse model of Staphylococcus aureus peritoneal infection. Infect Immun 80:3721–32.

12. Kielian T, Cheung A, Hickey WF. 2001. Diminished virulence of an alpha-toxin mutant of Staphylococcus aureus in experimental brain abscesses. Infect Immun 69:6902–11.

13. Hua L, Hilliard JJ, Shi Y, Tkaczyk C, Cheng LI, Yu X, Datta V, Ren S, Feng H, Zinsou R, Keller A, O’Day T, Du Q, Cheng L, Damschroder M, Robbie G, Suzich J, Stover CK, Sellman BR. 2014. Assessment of an anti-alpha-toxin monoclonal antibody for prevention and treatment of Staphylococcus aureus-induced pneumonia. Antimicrob Agents Chemother 58:1108–17.

14. Diep BA, Hilliard JJ, L. VT, Tkaczyk C, Le HN, Tran VG, Rao RL, Dip EC, Pereira-Franchi EP, Cha P, Jacobson S, Broome R, Cheng LI, Weiss W, Prokai L, Nguyen V, Stover CK, Sellman BR. 2017. Targeting Alpha Toxin To Mitigate Its Lethal Toxicity in Ferret and Rabbit Models of Staphylococcus aureus Necrotizing Pneumonia. Antimicrob Agents Chemother 61.

15. Sampedro GR, DeDent AC, Becker RE, Berube BJ, Gebhardt MJ, Cao H, Bubeck Wardenburg J. 2014. Targeting Staphylococcus aureus alpha-toxin as a novel approach to reduce severity of recurrent skin and soft-tissue infections. J Infect Dis 210:1012–8.

16. Le VT, Tkaczyk C, Chau S, Rao RL, Dip EC, Pereira-Franchi EP, Cheng L, Lee S, Koelkebeck H, Hilliard JJ, Yu XQ, Datta V, Nguyen V, Weiss W, Prokai L, O’Day T, Stover CK, Sellman BR, Diep BA. 2016. Critical Role of Alpha-Toxin and Protective Effects of Its Neutralization by a Human Antibody in Acute Bacterial Skin and Skin Structure Infections. Antimicrob Agents Chemother 60:5640–8.

17. Adhikari RP, Ajao AO, Aman MJ, Karauzum H, Sarwar J, Lydecker AD, Johnson JK, Nguyen C, Chen WH, Roghmann MC. 2012. Lower antibody levels to Staphylococcus aureus exotoxins are associated with sepsis in hospitalized adults with invasive S. aureus infections. J Infect Dis 206:915–23.

18. Li S, Arvidson S, Mollby R. 1997. Variation in the agr-dependent expression of alpha-toxin and protein A among clinical isolates of Staphylococcus aureus from patients with septicaemia. FEMS Microbiol Lett 152:155–61.

19. Stulik L, Malafa S, Hudcova J, Rouha H, Henics BZ, Craven DE, Sonnevend AM, Nagy E. 2014. alpha-Hemolysin activity of methicillin-susceptible Staphylococcus aureus predicts ventilator-associated pneumonia. Am J Respir Crit Care Med 190:1139–48.

20. Soriano A, Marco F, Martinez JA, Pisos E, Almela M, Dimova VP, Alamo D, Ortega M, Lopez J, Mensa J. 2008. Influence of vancomycin minimum inhibitory concentration on the treatment of methicillin-resistant Staphylococcus aureus bacteremia. Clin Infect Dis 46:193–200.

21. Harris PA, Taylor R, Thielke R, Payne J, Gonzalez N, Conde JG. 2009. Research electronic data capture (REDCap)--a metadata-driven methodology and workflow process for providing translational research informatics support. J Biomed Inform 42:377–81.

22. Kennedy AD, Bubeck Wardenburg J, Gardner DJ, Long D, Whitney AR, Braughton KR, Schneewind O, DeLeo FR. 2010. Targeting of alpha-hemolysin by active or passive immunization decreases severity of USA300 skin infection in a mouse model. J Infect Dis 202:1050–8.

23. Bhakdi S, Tranum-Jensen J. 1991. Alpha-toxin of Staphylococcus aureus. Microbiol Rev 55:733–51.

24. Novick RP, Ross HF, Projan SJ, Kornblum J, Kreiswirth B, Moghazeh S. 1993. Synthesis of staphylococcal virulence factors is controlled by a regulatory RNA molecule. EMBO J 12:3967–75.

25. Sharma-Kuinkel BK, Wu Y, Tabor DE, Mok H, Sellman BR, Jenkins A, Yu L, Jafri HS, Rude TH, Ruffin F, Schell WA, Park LP, Yan Q, Thaden JT, Messina JA, Fowler VG, Jr., Esser MT. 2015. Characterization of alpha-toxin hla gene variants, alpha-toxin expression levels, and levels of antibody to alpha-toxin in hemodialysis and postsurgical patients with Staphylococcus aureus bacteremia. J Clin Microbiol 53:227–36.

26. Tavares A, Nielsen JB, Boye K, Rohde S, Paulo AC, Westh H, Schonning K, de Lencastre H, Miragaia M. 2014. Insights into alpha-hemolysin (Hla) evolution and expression among Staphylococcus aureus clones with hospital and community origin. PLoS One 9:e98634.

27. Gafter-Gvili A, Mansur N, Bivas A, Zemer-Wassercug N, Bishara J, Leibovici L, Paul M. 2011. Thrombocytopenia in Staphylococcus aureus bacteremia: risk factors and prognostic importance. Mayo Clin Proc 86:389–96.

28. Akca S, Haji-Michael P, de Mendonca A, Suter P, Levi M, Vincent JL. 2002. Time course of platelet counts in critically ill patients. Crit Care Med 30:753–6.

29. Vandijck DM, Blot SI, De Waele JJ, Hoste EA, Vandewoude KH, Decruyenaere JM. 2010. Thrombocytopenia and outcome in critically ill patients with bloodstream infection. Heart Lung 39:21–6.

30. Coia JE, Browning L, Haines L, Birkbeck TH, Platt DJ. 1992. Comparison of enterotoxins and haemolysins produced by methicillin-resistant (MRSA) and sensitive (MSSA) Staphylococcus aureus. J Med Microbiol 36:164–71.

31. Jacobsson G, Colque-Navarro P, Gustafsson E, Andersson R, Mollby R. 2010. Antibody responses in patients with invasive Staphylococcus aureus infections. Eur J Clin Microbiol Infect Dis 29:715–25.

32. Inagaki K, Lucar J, Blackshear C, Hobbs CV. 2019. Methicillin-susceptible and Methicillin-resistant Staphylococcus aureus Bacteremia: Nationwide Estimates of 30-Day Readmission, In-hospital Mortality, Length of Stay, and Cost in the United States. Clin Infect Dis 69:2112–2118.

33. Whitby M, McLaws ML, Berry G. 2001. Risk of death from methicillin-resistant Staphylococcus aureus bacteraemia: a meta-analysis. Med J Aust 175:264–7.

34. Cosgrove SE, Sakoulas G, Perencevich EN, Schwaber MJ, Karchmer AW, Carmeli Y. 2003. Comparison of mortality associated with methicillin-resistant and methicillinsusceptible Staphylococcus aureus bacteremia: a meta-analysis. Clin Infect Dis 36:53–9.

35. Cheung GY, Wang R, Khan BA, Sturdevant DE, Otto M. 2011. Role of the accessory gene regulator agr in community-associated methicillin-resistant Staphylococcus aureus pathogenesis. Infect Immun 79:1927–35.

36. Otto M. 2012. MRSA virulence and spread. Cell Microbiol 14:1513–21.

37. Chua KY, Monk IR, Lin YH, Seemann T, Tuck KL, Porter JL, Stepnell J, Coombs GW, Davies JK, Stinear TP, Howden BP. 2014. Hyperexpression of alpha-hemolysin explains enhanced virulence of sequence type 93 community-associated methicillin-resistant Staphylococcus aureus. BMC Microbiol 14:31.

38. Li M, Dai Y, Zhu Y, Fu CL, Tan VY, Wang Y, Wang X, Hong X, Liu Q, Li T, Qin J, Ma X, Fang J, Otto M. 2016. Virulence determinants associated with the Asian community-associated methicillin-resistant Staphylococcus aureus lineage ST59. Sci Rep 6:27899.

39. Yamaki J, Synold T, Wong-Beringer A. 2013. Tigecycline induction of phenol-soluble modulins by invasive methicillin-resistant Staphylococcus aureus strains. Antimicrob Agents Chemother 57:4562–5.

40. Berube BJ, Sampedro GR, Otto M, Bubeck Wardenburg J. 2014. The psmalpha locus regulates production of Staphylococcus aureus alpha-toxin during infection. Infect Immun 82:3350–8.

41. Dinges MM, Orwin PM, Schlievert PM. 2000. Exotoxins of Staphylococcus aureus. Clin Microbiol Rev 13:16-34, table of contents.

42. Oliveira D, Borges A, Simoes M. 2018. Staphylococcus aureus Toxins and Their Molecular Activity in Infectious Diseases. Toxin (Basel) 10.

